# An Integrated Biophysical Fragment Screening Approach Identifies Novel Binders of the CD28 Immune Receptor

**DOI:** 10.1101/2025.11.23.690058

**Authors:** Laura Calvo-Barreiro, Hossam Nada, Moustafa T. Gabr

## Abstract

CD28 is an essential costimulatory receptor required for full T cell activation, and its dysregulation contributes to multiple immune-mediated pathologies. Despite its central immunological role, CD28 remains largely unexplored as a target for small-molecule modulation, primarily due to the shallow and large interface of its ligand-binding site. Here, we applied a fragment-based high-throughput screening (HTS) strategy to identify low molecular weight chemotypes capable of engaging with human CD28. A 3,200-member library composed of structurally diverse fragments, enriched for scaffolds designed to target protein-protein interaction (PPI) interfaces, was screened in single-dose format using temperature-related intensity change (TRIC) technology, yielding 36 primary hits (1.13% hit rate). Follow-up surface plasmon resonance (SPR) validation confirmed two fragments as direct CD28 binders. Molecular docking analysis revealed a plausible binding orientation for PPIF3 within the CD28 extracellular domain, suggesting potential interaction hotspots that may be exploited during future optimization. Together, these findings provide the first demonstration that fragment-based screening can successfully identify chemotypes capable of engaging with the CD28 PPI interface. This work establishes a scalable, biophysics-driven workflow for CD28 ligand discovery and lays the foundation for subsequent hit-to-lead development of small molecule CD28 modulators.

## 1. Introduction

The costimulatory receptor CD28 is a central regulator of T-cell activation, survival, and homeostasis. By integrating signals from antigen-presenting cells through binding to its ligands (CD80/CD86), CD28 ensures proper immune priming and effector responses. Dysregulated CD28 signaling contributes to multiple immune-mediated pathologies, including autoimmunity, transplant rejection, and aberrant T-cell activation in cancer^1^. Given its pivotal immunological role and tractable extracellular domain, CD28 represents an attractive therapeutic target for selective modulation of T-cell function. Despite this importance, CD28 has remained largely underexplored as a small-molecule target due to the inherent challenges of its ligand-binding surface. Structural studies have shown that CD28 recognizes CD80/CD86 through a shallow, relatively flat protein-protein interaction (PPI) interface, which lacks deep pockets commonly exploited by drug-like molecules^1^. As a result, conventional screening approaches often struggle to identify tractable chemical matter for this receptor.

Over the past decade, small-molecule approaches have gained momentum as complementary strategies to biologics for targeting immune checkpoints^2^. Traditional high-throughput screening (HTS) can identify potent inhibitors or modulators, but the complexity and low druggability of PPIs make the discovery of novel chemotypes challenging. Fragment-based drug discovery (FBDD) has emerged as a powerful alternative, enabling the detection of low-molecular-weight ligands that engage shallow binding surfaces with high ligand efficiency. These fragment hits can be elaborated into more potent, selective scaffolds using medicinal chemistry and structure-based optimization^3^. FBDD is particularly well suited for receptors like CD28 because fragments are capable of probing sparse or topologically feature-poor surfaces, revealing interaction hot spots that may remain undetected by traditional screening chemotypes. However, the identification of fragment binders at challenging PPI interfaces requires biophysical methods with the sensitivity to detect weak interactions.

In this work, we applied a fragment-library screening strategy to identify novel small-molecule chemotypes targeting human CD28. A custom library of 3,200 structurally diverse fragments was screened using HTS, yielding 36 primary hits. Surface plasmon resonance (SPR) validation confirmed two fragments as direct CD28 binders, and one of them displayed a measurable affinity with a K_D_ of 395.33 ± 63.31 μM. Finally, molecular docking analysis provided initial insights into the putative binding pose and potential interaction hotspots. Together, these results demonstrate that even a shallow and historically difficult immune receptor such as CD28 can be engaged by low-molecular-weight fragments when leveraging sensitive biophysical detection methods. This study establishes a foundation for future hit-to-lead optimization and provides a scalable strategy for discovering small-molecule modulators of the CD28 immune checkpoint.

## 2. Methods

### 2.1. Screened Library

A subset of 3,200 compounds from the PPI Fragment Library (Cat. # PPIF-3600, Enamine, Kyiv, Ukraine) was selected as the compound library targeting the human CD28 protein. All compounds were contained in 384-well plates and dissolved in DMSO at a concentration equal to 10 mM. Libraries were stored at -80°C upon arrival and until use.

Following HTS, newly lyophilized material for each selected primary hit was obtained from Enamine. These compounds were reconstituted in DMSO at a concentration of 50 mM and stored at -30 °C until use.

### 2.2. Dianthus High-Throughput Affinity Screening

Affinity screening was performed using the Dianthus NT.23 Pico platform (NanoTemper Technologies, München, Germany) as previously described^4^. Briefly, His-tagged human CD28 protein (Cat. # CD8-H52H3, Acro Biosystems, Newark, DE, USA) was labeled with Monolith His-Tag Labeling Kit RED-tris-NTA 2^nd^ Generation (Cat. # MO-L018, NanoTemper Technologies) and incubated with test compounds or controls. Fluorescence changes were measured via temperature-related intensity change (TRIC) to assess binding when compared to the negative control. Compounds showing interaction with the dye alone were excluded. Confirmed binders were selected for confirmatory testing using orthogonal validation by SPR.

### 2.3. Biacore™ 8K Single-Dose Screening

Single-dose binding assays (200 µM) were performed using the Biacore™ 8K system (Cytiva, Marlborough, MA, USA) at 25 °C, following previously published protocol and data analysis^5^. Biotinylated human CD28 (Cat. # CD8-H82E5, Acro Biosystems) was immobilized using the Biotin CAPture Kit (Cat. # BR100669, Cytiva), and candidate compounds were injected in 1x PBS-P buffer with 2% DMSO. Anti-CD28 antibody served as a positive control, and solvent corrections were applied to account for DMSO-related refractive index changes. Data acquisition and analysis were conducted using Biacore™ 8K Control and Insight Evaluation Software, respectively, to identify and rank primary hits.

### 2.4. SPR-Based Binding Affinity Screening

Binding kinetics of selected hits were evaluated using a multi-cycle assay on the Biacore™ 8K system (Cytiva) at 25 °C, following previously published procedures^5^. Briefly, biotinylated CD28 was immobilized using the Biotin CAPture Kit, and analytes were injected at ascending concentrations (ranging from 43.9 µM up to 500 µM) in 1x PBS-P buffer with 2% DMSO. Solvent corrections and anti-CD28 antibody control were included to ensure data quality. Sensorgrams were analyzed using Biacore™ Insight Evaluation Software to determine steady-state affinities.

### 2.5. Molecular Docking

The molecular docking employed in this study followed the same computational methodology as previously described in our recently published work^5, 6^.

## 3. Results and Discussion

### 3.1. Fragment Screening of CD28 Using TRIC Technology

Building on the optimized TRIC workflow previously developed for CD28^4^, we applied the same assay design and criteria to screen a subset of the Enamine PPI Fragment Library^7^. TRIC technology enables the screening of low molecular-weight (MW) compounds thanks to its mass-independent technology in a solution-based 384-well plate format. This combination of features allows our biophysical screen to closely approximate physiological conditions, while avoiding the loss of sensitivity that can occur on other platforms when working with very small molecules (MW ranging between 190 Da and 460 Da). This screening collection contains 3,200 structurally diverse fragments specifically engineered to explore PPI interfaces and enriched in substructures known to interact with hot-spot residues. Because CD28 engages with its ligands (CD80/CD86) through a shallow, relatively flat interface, this library provided an appropriate source of low MW chemotypes with physicochemical properties (balanced polarity, scaffold diversity, and PPI-mimicking features) well suited for detecting weak but specific binders^7^.

Using freshly labeled CD28 protein and the optimized PBS-based buffer, we screened all 3,200 fragments at a final concentration of 200 µM. Each plate included two reference columns containing only assay buffer supplemented with DMSO, serving as negative controls (Figure 1, Primary Screening Output). As in our previously validated HTS workflow, a single F_norm_ value was obtained for each fragment and compared to the distribution of reference wells to calculate the ΔF_norm_ response. Compounds exceeding three times the standard deviation of the negative control were initially flagged as potential binders. To reduce false positives and maintain consistency with the criteria applied in our earlier screening campaign, we imposed an additional ΔF_norm_ threshold of 10 units (approximately 50% of the positive-control amplitude)^4^. Additionally, control experiments allowed us to identify fragments showing autofluorescence, quenching effects, or signal anomalies in the TRIC trace and, consequently, were removed from further consideration. After triage, 36 fragments remained as primary hits and were cherry-picked for follow-up testing (Table 1; Figure 1, Primary Screening Output).

**Table 1.** Primary fragment hits identified from the TRIC-based high-throughput screen and validated using SPR. The table summarizes biophysical parameters for all compounds progressing from the TRIC primary screen to SPR follow-up validation. For each analyte, the molecular weight (MW), analyte binding level (measured at the end of the SPR association phase), immobilized ligand level, calculated theoretical R_max_, and resulting Level of Occupancy (LO) are reported. Compounds were advanced to affinity characterization if they displayed an LO between 50–100% and were not flagged as non-specific binders (NSB). All measurements were obtained using biotinylated human CD28 ectodomain (70 kDa) as the immobilized ligand. Abbreviations: LO, level of occupancy; MW, molecular weight; NSB, non-specific binder; RU, response units.

| Analyte | Analyte MW (Da) | Analyte Binding Level (RU) | Immobilized Ligand Level (RU)* | $R_{max}$ (RU) | LO (%) | Curve Marker |
| --- | --- | --- | --- | --- | --- | --- |
| PPIF1 | 309.4 | 12.1 | 1646.1 | 14.55 | 82.95 |  |
| PPIF2 | 279.2 | 10.4 | 1646.1 | 13.13 | 78.93 | NSB |
| PPIF3 | 394.5 | 11.6 | 1646.1 | 18.55 | 62.37 |  |
| PPIF4 | 259.3 | 1.4 | 1682.8 | 12.47 | 11.16 |  |
| PPIF5 | 354.8 | 1.1 | 1646.1 | 16.69 | 6.577 | NSB |
| PPIF6 | 337.3 | 0.6 | 1703.3 | 16.41 | 3.844 |  |
| PPIF7 | 361.8 | 0.2 | 1646.1 | 17.02 | 1.188 |  |
| PPIF8 | 371.5 | -0.3 | 1703.3 | 18.08 | -1.61 |  |
| PPIF9 | 369.5 | -0.3 | 1646.1 | 17.38 | -1.858 |  |
| PPIF10 | 359.4 | -0.8 | 1703.3 | 17.49 | -4.417 |  |
| PPIF11 | 372.8 | -0.8 | 1703.3 | 18.14 | -4.584 |  |
| PPIF12 | 344.8 | -0.8 | 1682.8 | 16.58 | -4.76 |  |
| PPIF13 | 278.8 | -0.8 | 1682.8 | 13.4 | -5.806 |  |
| PPIF14 | 373.4 | -1.1 | 1703.3 | 18.17 | -5.816 |  |
| PPIF15 | 342.4 | -1.3 | 1646.1 | 16.1 | -8.323 |  |
| PPIF16 | 376.3 | -1.8 | 1682.8 | 18.09 | -9.751 |  |
| PPIF17 | 364.9 | -1.7 | 1646.1 | 17.16 | -9.894 |  |
| PPIF18 | 265.7 | -1.3 | 1703.3 | 12.93 | -10.16 |  |
| PPIF19 | 352.4 | -2.1 | 1682.8 | 16.94 | -12.52 |  |
| PPIF20 | 359.8 | -2.8 | 1703.3 | 17.51 | -16.04 |  |
| PPIF21 | 379.5 | -3.4 | 1703.3 | 18.47 | -18.52 |  |
| PPIF22 | 349.4 | -3.3 | 1682.8 | 16.8 | -19.82 |  |
| PPIF23 | 231.3 | -2.4 | 1703.3 | 11.26 | -21.28 |  |
| PPIF24 | 397.4 | -4.2 | 1646.1 | 18.69 | -22.58 |  |
| PPIF25 | 312.4 | -5 | 1682.8 | 15.02 | -33.21 |  |
| PPIF26 | 349.4 | -6 | 1682.8 | 16.8 | -35.6 | NSB |
| PPIF27 | 348.4 | -6.5 | 1703.3 | 16.96 | -38.37 |  |
| PPIF28 | 385.5 | -7.3 | 1646.1 | 18.13 | -40.28 | NSB |
| PPIF29 | 347.4 | -8 | 1703.3 | 16.91 | -47.27 | NSB |
| PPIF30 | 362.4 | -9.2 | 1646.1 | 17.04 | -54.05 | NSB |
| PPIF31 | 345.4 | -12.8 | 1682.8 | 16.61 | -76.95 | NSB |
| PPIF32 | 348.8 | -17 | 1703.3 | 16.97 | -99.91 |  |
| PPIF33 | 398.5 | -20.6 | 1682.8 | 19.16 | -107.5 | NSB |
| PPIF34 | 367.4 | -24.5 | 1682.8 | 17.66 | -138.6 |  |
| PPIF35 | 349.4 | -23.8 | 1682.8 | 16.8 | -141.6 | NSB |

**Figure 1.**
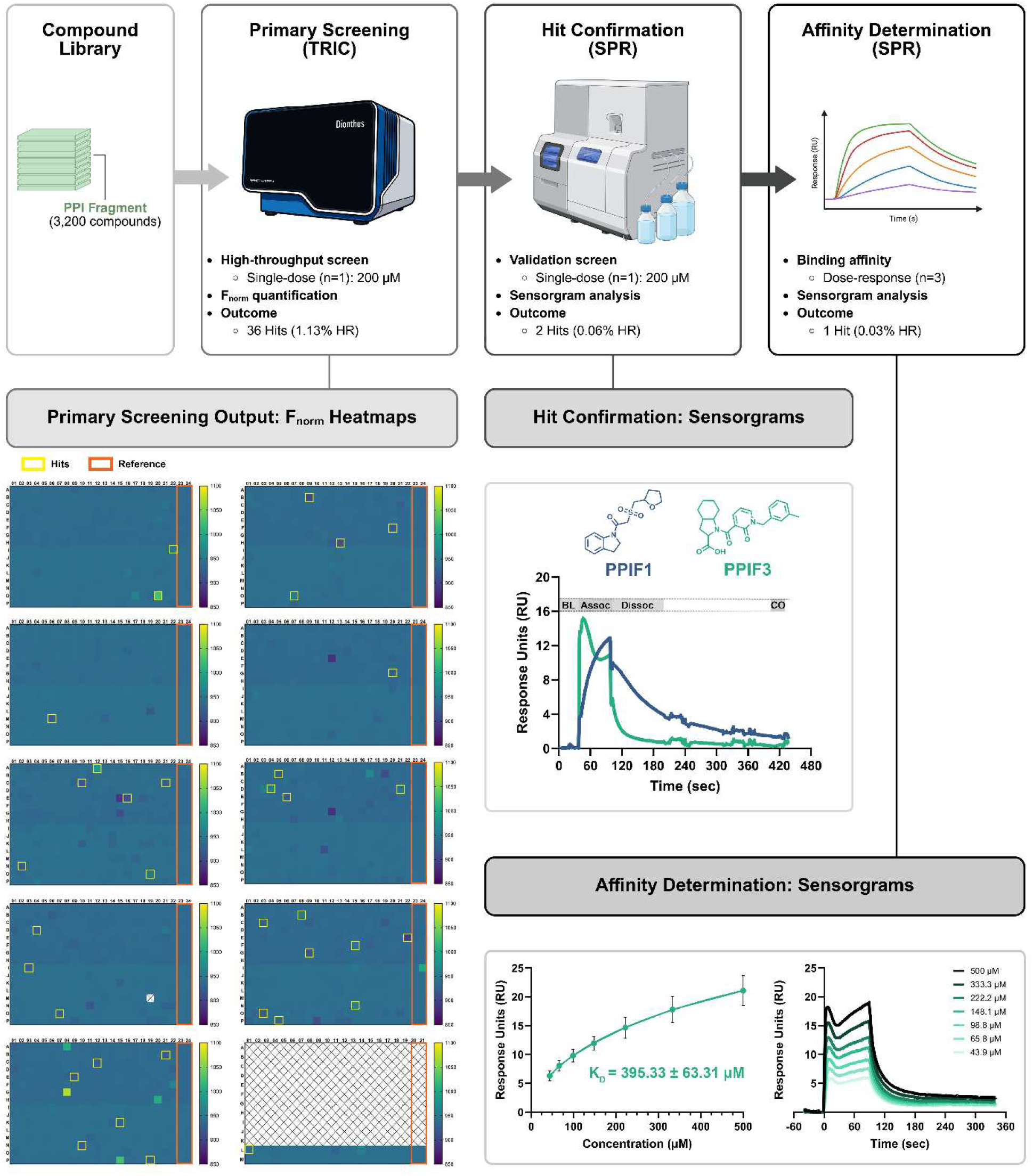
Workflow and results of the primary screening, hit confirmation, and affinity determination for CD28-targeted PPI fragments. Schematic overview of the screening funnel integrating TRIC-based primary screening with SPR-based hit validation and affinity characterization. A 3,200-member PPI Fragment library was screened in single-dose format (200 µM), yielding 36 primary hits (1.13% HR) based on F_norm_ output. 384-well plate heatmaps illustrate F_norm_ distributions and highlight wells exceeding hit thresholds (yellow boxes) and reference wells included as experimental negative controls (orange wells). Selected primary hits were subjected to SPR validation using a single-dose injection (200 µM), resulting in two confirmed binders (0.06% HR) with clear association and dissociation phases. Representative sensorgrams for the two validated hits, PPIF1 and PPIF3, are shown. Following a stable baseline (BL), each 200 µM fragment solution was injected over the sensor surface for 60 sec (association phase, Assoc), and the resulting response (RU) relative to baseline was recorded. A 100 sec dissociation phase (Dissoc) allowed disengagement from the active flow cell, after which a carryover control (CO) step was performed by flowing additional running buffer over both flow cells. Subsequent dose-response SPR analysis identified one fragment with quantifiable binding affinity (K_D_ = 395.33 ± 63.31 µM; 0.03% overall HR). Fitted steady-state curves (three independent experiments, mean ± SD) and representative sensorgrams (one independent experiment) are shown. Abbreviations: HR: Hit Rate, PPI: Protein-Protein Interaction, SPR: Surface Plasmon Resonance, TRIC: Temperature-Related Intensity Change.

### 3.2. Biophysical Validation of Hits Using SPR

Following TRIC-based hit selection, commercially available primary fragments (35 out of the initial 36 candidates) were evaluated using SPR to assess direct binding to immobilized CD28 (Table 1). We adopted an SPR workflow aligned with the previously published CD28 HTS platform, screening fragments based on level of occupancy (LO), response magnitude, and dissociation kinetics^5^. Hits were considered validated only when they achieved LO > 50%, displayed clean association/dissociation profiles, and passed all specificity filters, including absence of non-specific binding to the reference flow-cell and lack of drift or non-dissociating behavior. This strict gate mirrors the validated criteria reported in the original study and ensures high confidence in true positives.

Using this orthogonal validation approach, two fragments exhibited measurable binding responses at 200 µM in a single-dose experiment (Figure 1, Hit Confirmation). For one of them, PPIF3, a full concentration series (ranging 500 to 43.9 µM) enabled determination of an affinity constant, yielding a K_D_ of 395.33 ± 63.31 µM, consistent with fragment-like binding (Figure 1, Affinity Determination)^8^. The second hit, PPIF1, showed clear binding but did not reach saturation within the tested concentration range, preventing reliable K_D_ estimation. Overall, SPR validation not only confirmed direct binding but also provided kinetic insights that strengthened prioritization of the selected fragments for downstream characterization.

### 3.3. Docking Analysis to Explore Binding Interactions

The costimulatory receptor CD28 and its inhibitory counterpart CTLA-4 engage B7 family ligands (CD80/CD86) through a shared binding mechanism. Crystallographic analyses have shown that their primary ligand-binding region resides within a conserved segment encompassing residues 99-104^9^. Mutagenesis studies further confirmed the functional importance of this site, as substitutions within this sequence cause a marked (>90%) reduction in ligand-binding affinity. Despite its critical role in receptor-ligand recognition, the CD28 binding interface poses major challenges for small-molecule drug design. Structural analysis of the CD28-CD80 complex (PDB ID: 1YJD) reveals a relatively flat and extended interaction surface, lacking the deep, well-defined pockets typically required for high-affinity small-molecule binding. Such shallow protein-protein interfaces are inherently difficult to target with drug-like compounds because they offer limited opportunities for specific and stable molecular interactions^5^.

To gain insight into the molecular mechanisms underlying the biological activity of PPIF3, we performed molecular docking studies to characterize its interaction with the CD28 receptor. The docking results revealed that PPIF3 adopts a distinct binding orientation within the CD28 ligand-binding pocket, differing spatially from previously reported ligands (Figure 2A-B)^5^. Detailed analysis of the docking pose indicated that PPIF3 forms key hydrogen bonds with Lys95, Lys109, and Asp106, which are known to be critical residues for ligand recognition. In addition, Phe93 contributes to complex stabilization through hydrophobic π-π stacking interactions with the aromatic core of PPIF3, collectively suggesting a well-defined and energetically favorable binding mode (Figure 2B).

**Figure 2.**
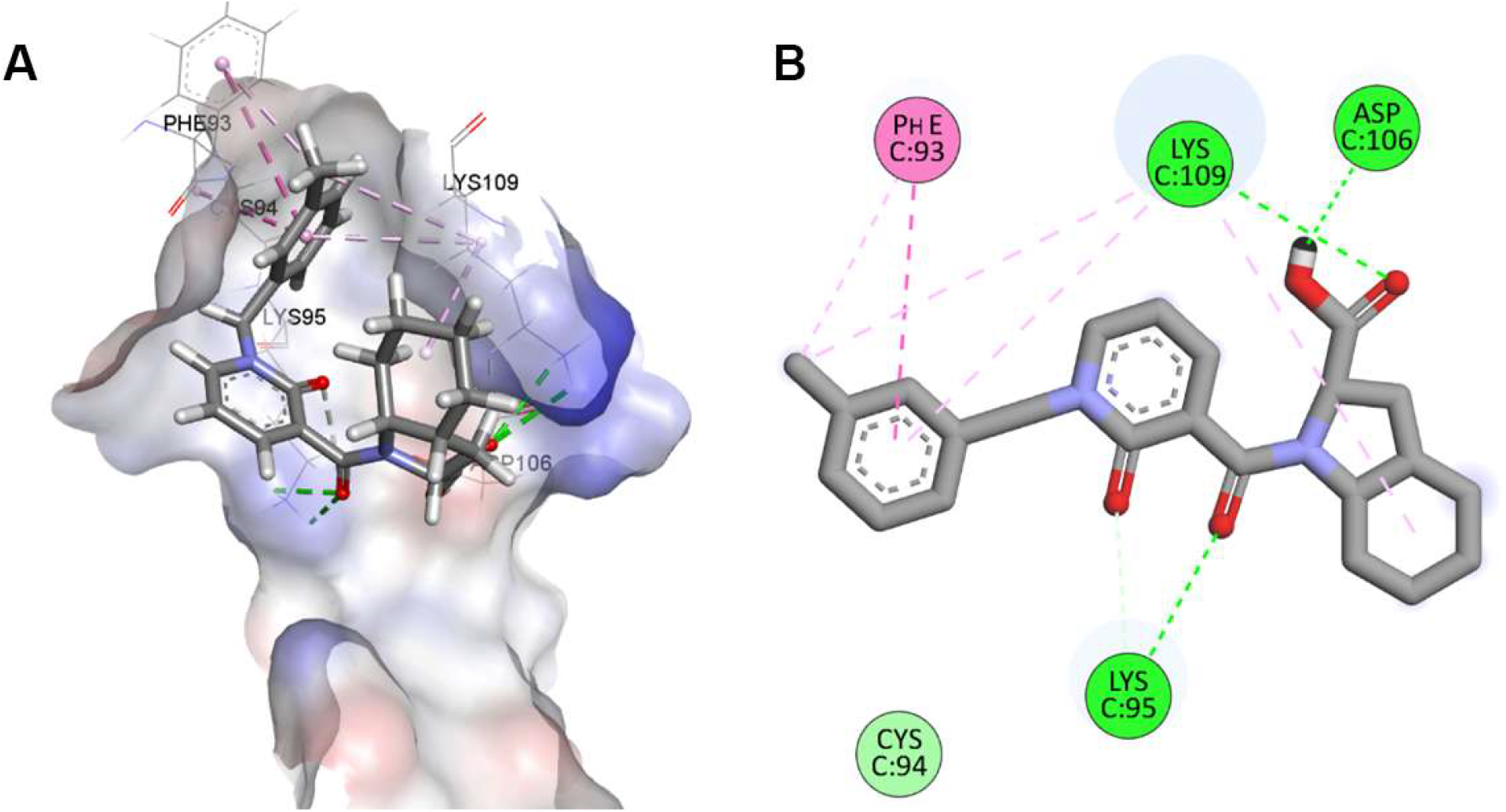
Molecular docking analysis of PPIF3 binding to CD28. (A) Three-dimensional representation of PPIF3 bound to the CD28 receptor. (B) Two-dimensional interaction diagram of PPIF3 with CD28

Several previously reported CD28-targeted small molecules, including those identified through virtual screening, mass spectrometry, and TRIC-based HTS workflows,^4-6,10^ belong predominantly to higher molecular weight drug-like scaffolds. These chemotypes generally rely on multi-point interactions distributed across the relatively flat CD28-B7 interface. Because such compounds occupy a larger surface footprint, their engagement often depends on combining hydrophobic surface complementarity with extended π-systems or hydrogen-bonding networks spanning multiple secondary structural elements. This recognition pattern aligns with expectations for PPI-directed inhibitors, which often require considerable surface area coverage to compensate for the absence of deep, pocket-like cavities.

In contrast, the fragment hit identified here, PPIF3, is substantially smaller and exhibits a more compact pharmacophore. Our docking analysis indicates that PPIF3 does not mimic the canonical binding profiles observed for previously described small molecules. Instead, PPIF3 appears to localize within a more restricted microenvironment of the CD28 interface, forming a focused cluster of interactions with Lys95, Asp106, Lys109, and Phe93. These residues are known to lie within or adjacent to the CD80/CD86 contact region, yet the specific spatial configuration adopted by PPIF3 differs from the surface-spanning arrangements formed by larger ligands. This suggests that PPIF3 is not merely a downsized version of existing chemotypes, but rather a fragment that exploits an orthogonal interaction motif within the CD28 extracellular domain.

The identification of such an alternative binding orientation is noteworthy for two reasons. First, it demonstrates that the CD28 interface, previously considered poorly druggable due to its shallow topology, contains localized hot spots that can support fragment-sized interactions. Second, it demonstrates that fragment-based screening can identify chemotypes that engage the target through interaction modes not accessible to standard HTS libraries. Whereas larger molecules typically depend on broad surface engagement, fragments can bind by anchoring into subtle topological depressions, forming high-ligand-efficiency interactions that may remain undetected by larger scaffolds.

Although direct comparison between fragment hits and drug-like CD28 ligands must be interpreted cautiously, given differences in affinity, size, and screening modality, the distinct binding mode observed for PPIF3 underscores the complementary nature of fragment-based approaches. Fragments such as PPIF3 can reveal previously unappreciated binding vectors, identify incipient pockets or transient subpockets, and provide medicinal chemistry starting points that are chemically tractable and structurally divergent from prior scaffolds. Collectively, these findings indicate that biophysical fragment screening can expand the accessible chemical space for CD28 modulation and may ultimately enable the rational evolution of fragments into higher-affinity ligands with unique mechanisms of action.

## 4. Conclusions

We have conducted the largest HTS to date using fragment-based approaches targeting the CD28 immune receptor with biophysical techniques. From a 3,200-fragment library, we identified and validated an efficient fragment that binds to the protein surface. The overall low hit rate of 0.03% underscores the challenges inherent to this class of targets which has also been observed in previous campaigns^4-6,10^. Thus, our findings highlight both the structural difficulties and the opportunities in modulating the CD28-B7 costimulatory axis with small molecules.

Although the CD28 ligand-binding interface is shallow and lacks well-defined pockets, our molecular docking studies demonstrate that selective fragment engagement is achievable. PPIF3 adopts a distinct and energetically favorable binding mode within the CD28 interface. Its ability to form stabilizing hydrogen bonds with key recognition residues (Lys95, Asp106, Lys109) and hydrophobic π-π interactions with Phe93 supports a plausible molecular mechanism for its observed micromolar activity. These results provide a structural rationale for fragment binding at this challenging interface and lay the foundation for structure-guided optimization toward higher-affinity CD28 modulators.

## Acknowledgments

This work was supported by the National Institute of Diabetes and Digestive and Kidney Diseases (NIDDK) under grant number R01DK137299. We would like to thank the Fisher Drug Discovery Resource Center of Rockefeller University (RRID:SCR_020985) for providing access to the Nanotemper Dianthus NT.23 Pico and Cytiva Biacore 8K instruments.

